# Smartwatch-based prediction of single-stride and stride-to-stride gait outcomes using regression-based machine learning

**DOI:** 10.1101/2023.01.30.526246

**Authors:** Christopher A. Bailey, Alexandre Mir-Orefice, Thomas K. Uchida, Julie Nantel, Ryan B. Graham

## Abstract

Spatiotemporal variability during gait is linked to fall risk and could be monitored using wearable sensors. Although many users prefer wrist-worn sensors, most applications position at other sites. We developed and evaluated an application using a consumer-grade smartwatch inertial measurement unit (IMU). Young adults (N = 41) completed seven-minute conditions of treadmill gait at three different speeds. Single-stride outcomes (stride time, length, width, and speed) and spatiotemporal variability (coefficient of variation of each single-stride outcome) were recorded using an optoelectronic system, while 232 single- and multi-stride IMU metrics were recorded using an Apple Watch Series 5. These metrics were input to train linear, ridge, support vector machine (SVM), random forest, and extreme gradient boosting (xGB) models of each spatiotemporal outcome. We conducted Model × Condition ANOVAs to explore model sensitivity to speed-related responses. xGB models were best for single-stride outcomes (relative mean absolute error [% error]: 7–11%; intraclass correlation coefficient [ICC_2,1_]: 0.60–0.86) and SVM models were best for spatiotemporal variability (% error: 18–22%; ICC_2,1_ = 0.47–0.64). Spatiotemporal changes with speed were captured by these models (Condition: p < 0.00625). Results support the feasibility of monitoring multi-stride spatiotemporal parameters using a smartwatch IMU and machine learning.

## 3. Introduction

Walking-related falls are linked to the natural motor fluctuations that exist from stride to stride ^16^. Gait investigations have found that older individuals with higher stride time variability ^17^, higher stride length variability ^26^, and very low or very high step width variability ^6^ have higher fall risk. The strong link between spatiotemporal variability in gait and fall risk calls for real-world evaluation methods that enable and support community-based monitoring.

Spatiotemporal gait evaluations have traditionally relied on in-lab or in-clinic methods. These include using marker-based optoelectronic motion capture ^46^, video-based optoelectronic motion capture ^23,39^, and pressure mats ^4^. These approaches, while considered valid and reliable for spatiotemporal analyses, are expensive (optoelectronic systems), require time-consuming measurement (marker-based optoelectronic systems), and restrict the area in which gait can be measured (all approaches listed above). A large motion capture volume, beyond that available from these approaches, is needed to capture 50–150 continuous strides to reliably measure spatiotemporal variability during overground walking ^24,37^. Thus, alternative methods are needed for inexpensive, continuous, and large-volume evaluations that are required in community settings.

Wearable devices may meet these needs, with two potential options being instrumented insoles and body-worn inertial measurement units (IMUs) ^7^. Instrumented insoles contain embedded pressure sensors and/or IMUs that directly measure temporal gait features via foot contact events ^40^, producing highly accurate detection of heel strikes, toe offs, and step count ^3,10,30^. Several open challenges exist in the use of instrumented insoles, including user comfort, compatibility in different footwear, and usability in different environmental conditions ^40^. An alternative option to insoles are body-worn IMUs, which have been used to calculate spatial and temporal gait features ^2,33,35,36,41,44^. Using a single foot IMU, Rebula et al. ^35^ developed an algorithm combining stride segmentation, drift correction, and foot trajectory formation that, relative to optoelectronic motion capture, predicted stride length and duration with a 1% difference and stride-to-stride length and width variability with a 4% difference. Washabaugh and colleagues ^44^ used foot and ankle IMUs, finding that spatiotemporal gait parameters were measured better by the foot configuration, were concurrently valid relative to an instrumented treadmill, and were repeatable between days. Using a single trunk IMU, de Ridder and colleagues ^36^ demonstrated validity and intra-day reliability in measuring gait speed, cadence, stride length, and stride time. IMUs can also be instrumented on multiple lower limb segments and combined with biomechanical constraints to model foot trajectories and extract spatiotemporal parameters ^2,41^. A major challenge with each of these IMU approaches is that the sensors are not positioned on locations preferred by most individuals, as exemplified in a recent investigation of persons with Parkinson’s disease ^32^, questioning the feasibility of these IMU positions for long-term continuous monitoring.

As a wrist-based IMU has been found to be preferred as a single-sensor solution ^32^, users may be more compliant to long-term monitoring with a smartwatch approach. Smartwatches are widely available and continue to grow in popularity as a smart wearable device due, in part, to perceived usefulness, enjoyment, and ease of use ^31^. In conjunction with machine learning techniques, wrist-based IMUs have been used for gait recognition ^21^, freezing of gait detection ^28,32^, fall detection ^43^, and spatiotemporal feature estimation ^12^. For example, Erdem and colleagues ^12^ used regression-based machine learning on linear acceleration and angular velocity features of smartwatches worn on both wrists to predict step length, swing time, and stance time to within 5.3 cm, 0.05 s, and 0.09 s, respectively. However, to the best of our knowledge, no application currently exists to predict spatiotemporal variability during gait using a single smartwatch, which is the most natural use case for this technology. Our objective was to develop and evaluate the accuracy of this application, using regression-based machine learning on features engineered from a single smartwatch IMU.

## 4. Materials and Methods

### 4.1 Participants

Young healthy adults (N = 41, 22 females; age: 25 ± 3 years, 19–31 years) were recruited to the study from the Ottawa, Canada region as a convenience sample. Participants were free from musculoskeletal injuries in the preceding six months and from known chronic neurological/orthopaedic disorders. All participants provided their written informed consent to the study. The study followed the Declaration of Helsinki and was approved by the University of Ottawa Research Ethics Board (H-01-21-6261).

### 4.2 Procedure

Each participant was instrumented for motion capture with an optoelectronic system and with a smartwatch. The optoelectronic system comprised 11 passive infrared cameras (Vantage, Vicon, Oxford, UK) and spherical retroreflective markers. Calibration and tracking markers were placed on each participant’s body according to a full-body marker set for gait ^2,34^ with rigid-body clusters of four markers on the trunk, arms, forearms, thighs, and shanks. Marker positions were sampled at 60 Hz using motion capture software (Nexus 2.11, Vicon, Oxford, UK). Each participant wore a smartwatch on their left wrist (Apple Watch Series 5, Apple Inc., Cupertino, USA). Tridimensional gravity-corrected linear accelerations (i.e. free accelerations), raw linear accelerations (i.e. raw accelerations), angular velocities, and orientations (Euler angles) were accessed from the Apple Core Motion API, via the HemiPhysioData app on watchOS, and were sampled at a requested frequency of 40 Hz.

Following a static calibration in standing pose, calibration markers were removed and tracking markers and smartwatch IMU data were sampled while the participant walked on a treadmill (Horizon Fitness, WI, USA). Preferred gait speed was identified according to the procedure of Dingwell and Marin ^11^. After establishing preferred gait speed, the participant completed three randomized speed conditions: preferred speed (Preferred), 70% of preferred speed (Slow), and 130% of preferred speed (Fast). Gait speed alters spatiotemporal variability ^22^, so the speed conditions were a method of exploring the sensitivity of the smartwatch IMU application. Each condition began with a vertical jump to synchronize the optoelectronic and smartwatch data streams and was followed by seven minutes of gait to record at least six minutes of consecutive and constant-speed strides. We determined during piloting that this duration was needed to confidently record a minimum of 150 steady-state strides for stable measurements of motor variability ^37^.

### 4.3 Data analysis

#### 4.3.1 Optoelectronic data processing

Using Vicon Nexus, marker trajectories were labelled, gap-filled with a Woltring spline ^45^, and low-pass filtered at 10 Hz. Processed marker trajectories were imported into OpenSim v4.2 ^38^ to simulate full-body motion ^2^. A generic skeletal model ^34^ was scaled to the participant by the anatomical marker positions in the static calibration and an inverse kinematic analysis was performed by minimizing the sum of weighted squared distance errors between pairs of experimental and model markers. Marker weights were manually selected to minimize root-mean-square error between marker pairs; marker weights were equal except for weights of double magnitude for markers on the acromion processes, anterior and posterior superior iliac spines, and lateral malleoli. Root-mean-square errors achieved following scaling and inverse kinematics were confirmed to be within the recommended range ^18^.

Modelled right calcaneus kinematics were subsequently analyzed in Matlab (R2021b, The MathWorks Inc., Natick, MA, USA). We partitioned individual right strides by identifying right heel strike events from the Euclidean norm of the right calcaneus linear velocity; events were identified as the local minima that followed a local maximum ^2^. These heel strike events and the modelled right calcaneus anteroposterior and mediolateral positions were then used to calculate stride time, length, width, and speed for the 200 strides following the initial 30 seconds of gait, during which time the participant was assumed to have reached steady state. We calculated the stride-to-stride coefficient of variation (CV) for each stride outcome to measure spatiotemporal variability.

#### 4.3.2 Smartwatch data processing

Raw inertial data from the smartwatch were sorted by timestamps, then resampled to 60 Hz to correct inconsistent time intervals between samples and to match the optoelectronic sampling frequency. Smartwatch data were synchronized to optoelectronic data by the peak Euclidean norm of the smartwatch linear acceleration and of the optoelectronic-modelled right calcaneus linear velocity upon landing from the vertical jump. Synchronized smartwatch data were then partitioned into individual strides by the optoelectronic-identified right heel strike events and time-normalized to 101 points per stride. The time-normalized stride and continuous formats of the smartwatch data series were retained for the same 200 strides as for the optoelectronic data. Smartwatch data included 16 IMU *Series*, representing the 4 *Inertial signals* (raw acceleration, free acceleration, angular velocity, Euler angle) and 4 *Components* of each Inertial signal (X, Y, Z, Euclidean norm of XYZ Components). From each Series, we extracted 16 *Metrics* of interest (Table 1). During data exploration for each Metric, we identified many extreme outliers for modulation and standard deviation (SD) of modulation, on the X, Y, and Z Components of each Inertial signal. These were produced by signal means approaching zero and were excluded from further analysis.

**Table 1.**
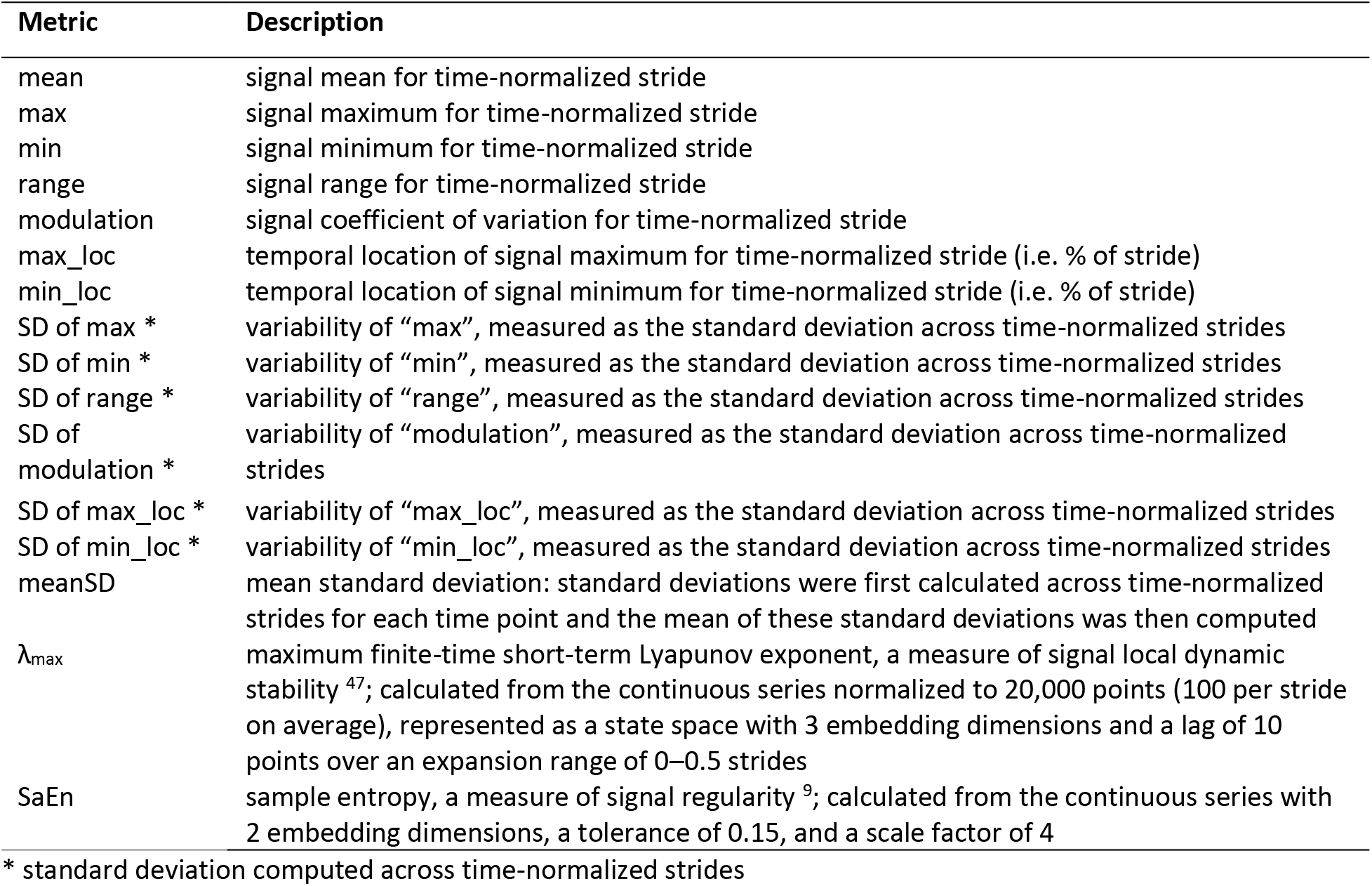
Smartwatch Metrics, calculated for each Component (X, Y, Z, Euclidean norm) of each Inertial signal (free acceleration, raw acceleration, angular velocity, Euler angle).

#### 4.3.3 Model development and evaluation

Before developing models of spatiotemporal variability, we first sought to make single-stride predictions as these outcomes are more common in prior applications (e.g., ^12^) and are available from consumer mobile devices using proprietary algorithms. Models of stride time, length, width, and speed were developed from smartwatch IMU Metrics taken from a single stride (mean, max, min, range, modulation, max_loc, min_loc; n = 112 Metrics). Accounting for excluded data (modulation of the X, Y, and Z Components, n = 12 Metrics), a total of 100 Metrics were used to develop single-stride models. Models of spatiotemporal variability were developed from all smartwatch IMU Metrics, including the across-stride means of single-stride smartwatch IMU Metrics and the multi-stride smartwatch IMU Metrics (n = 256 Metrics). After modulation and SD of modulation were excluded for the X, Y, and Z Components (n = 24 Metrics), this left 232 Metrics for developing models of spatiotemporal variability.

Smartwatch IMU Metrics and optoelectronic-measured spatiotemporal variability were available for 117 of 123 trials, producing 100 Metrics × 23,400 values for each single-stride outcome and 232 Metrics × 117 values for each spatiotemporal variability outcome. Principal component analysis was conducted on the smartwatch IMU inputs to each outcome group (single-stride outcomes, spatiotemporal variability) to extract the principal components that explained 90% of variation in each dataset. From each loading matrix, we weighted Metric loadings by the explained variance of each principal component, calculated the weighted mean of absolute loadings across principal components for each Metric, then identified the 20 Metrics with the highest weighted mean loadings. These top 20 Metrics were the regression model inputs. These data were split into 5 repeated folds, each consisting of model training (80%) and testing (20%) sets, where one set contained all trials for a participant. Repeated folds were used to evaluate model performance. For each repeated fold, we trained (i) linear, (ii) ridge, (iii) support vector machine (SVM), (iv) random forest (RF), and (v) extreme gradient boosting (xGB) regressors. Model inputs were z-scaled prior to training ridge and SVM regressors. Model training and testing were performed on a desktop computer containing one Radeon RX Vega GPU (8 Gb), one AMD Ryzen 7 2700x CPU (8 cores, 16 threads, 3.7 GHz), and 32 Gb of RAM. As SVM training exceeded the computational memory capacity when using all 200 strides to predict stride outcomes, we instead trained these models on only the first 50 strides per trial. We tuned hyperparameters of models (ii)–(v) on the training set by a randomized search of up to 100 iterations with 5-fold cross-validation, where 20% of the training set was used as a validation set (hyperparameter search ranges are provided in Appendix A and final hyperparameter-tuned models are provided in Appendix B). As xGB models tended to overfit, we terminated fitting early if test set accuracy did not improve for 10 consecutive epochs. Model accuracy was evaluated on the test set of each repeated fold by calculating the coefficient of determination (R^2^) and mean absolute error (MAE) between measured and predicted values.

### 4.4 Statistical analysis

Measured and predicted stride outcomes (across-stride mean for a given trial) and spatiotemporal variability were evaluated for consistency by computing two-way random intraclass correlation coefficients (ICC_2,1_), evaluated for agreement by computing bias and 95% limits of agreement (LOA_95%_), and visualized using Bland–Altman plots. ICC_2,1_ values less than 0.40, from 0.40 to 0.59, from 0.60 to 0.74, and greater than or equal to 0.75 were interpreted as poor, fair, good, and excellent consistency, respectively ^8^. Sensitivity of model predictions to within-subject changes in gait speed was evaluated by performing two-way repeated measures ANOVAs. ANOVAs tested for main effects and interactions of Condition (70%, 100%, 130% preferred speed) and Model (Measured, Linear, Ridge, SVM, RF, xGB). Greenhouse–Geisser corrections were applied when sphericity was violated; simple contrasts were made post-hoc relative to 100% preferred speed for Condition effects and relative to Measured values for Model effects. Statistical significance for all analyses was set at p < 0.00625 to adjust for the eight spatiotemporal outcomes (i.e. p < 0.050/8).

## 5. Results

### 5.1 Regression models

Table 2 lists the top 20 smartwatch IMU Metrics selected as inputs to the stride outcome and spatiotemporal variability models. Accuracy of stride outcome models is displayed in Figure 1. These predictions were generally best using xGB, with R^2^ = 0.61 ± 0.09 and MAE = 0.07 ± 0.01 s for stride time, R^2^ = 0.39 ± 0.20 and MAE = 0.13 ± 0.03 m for stride length, R^2^ = 0.46 ± 0.14 and MAE = 0.06 ± 0.01 m for stride width, and R^2^ = 0.69 ± 0.08 and MAE = 0.14 ± 0.02 m/s for stride speed. MAE values corresponded to relative errors of 7–11%. Spatiotemporal variability calculated from xGB-predicted stride outcomes was highly inaccurate, with MAE of 1.38–3.98% and relative errors of 63–272%.

**Table 2.** Smartwatch IMU metrics selected as regression inputs. Metrics were selected by the largest average absolute weighted load onto principal components extracted via principal component analysis.

|  | **Inertial signal** | **Axis** | **Metric** |
| --- | --- | --- | --- |
| **Stride outcome model** | Angular velocity | Z | range |
|  | Raw linear acceleration | Y | range |
|  | Angular velocity | Norm | mean |
|  | Raw linear acceleration | Z | range |
|  | Orientation | Y | range |
|  | Raw linear acceleration | X | mean |
|  | Angular velocity | Z | mean |
|  | Raw linear acceleration | Y | maximum |
|  | Raw linear acceleration | Y | minimum |
|  | Angular velocity | Z | maximum |
|  | Raw linear acceleration | Z | maximum |
|  | Raw linear acceleration | Norm | maximum |
|  | Raw linear acceleration | Norm | mean |
|  | Angular velocity | Norm | maximum |
|  | Angular velocity | Y | range |
|  | Angular velocity | Norm | range |
|  | Free linear acceleration | X | mean |
|  | Free linear acceleration | Norm | mean |
|  | Free linear acceleration | Norm | maximum |
|  | Angular velocity | Y | minimum |
| **Spatiotemporal variability model** | Raw linear acceleration | Y | range |
|  | Raw linear acceleration | Norm | mean |
|  | Raw linear acceleration | Y | meanSD |
|  | Angular velocity | Norm | meanSD |
|  | Raw linear acceleration | Y | maximum |
|  | Raw linear acceleration | Z | maximum |
|  | Free linear acceleration | Norm | maximum |
|  | Raw linear acceleration | Norm | maximum |
|  | Raw linear acceleration | X | maximum |
|  | Raw linear acceleration | Y | minimum |
|  | Angular velocity | Norm | mean |
|  | Raw linear acceleration | Norm | meanSD |
|  | Raw linear acceleration | Z | range |
|  | Orientation | Z | range |
|  | Raw linear acceleration | X | mean |
|  | Orientation | X | range |
|  | Angular velocity | Z | maximum |
|  | Angular velocity | Z | minimum |
|  | Orientation | Y | range |
|  | Angular velocity | X | meanSD |

*Norm*: Euclidean norm

**Figure 1.**
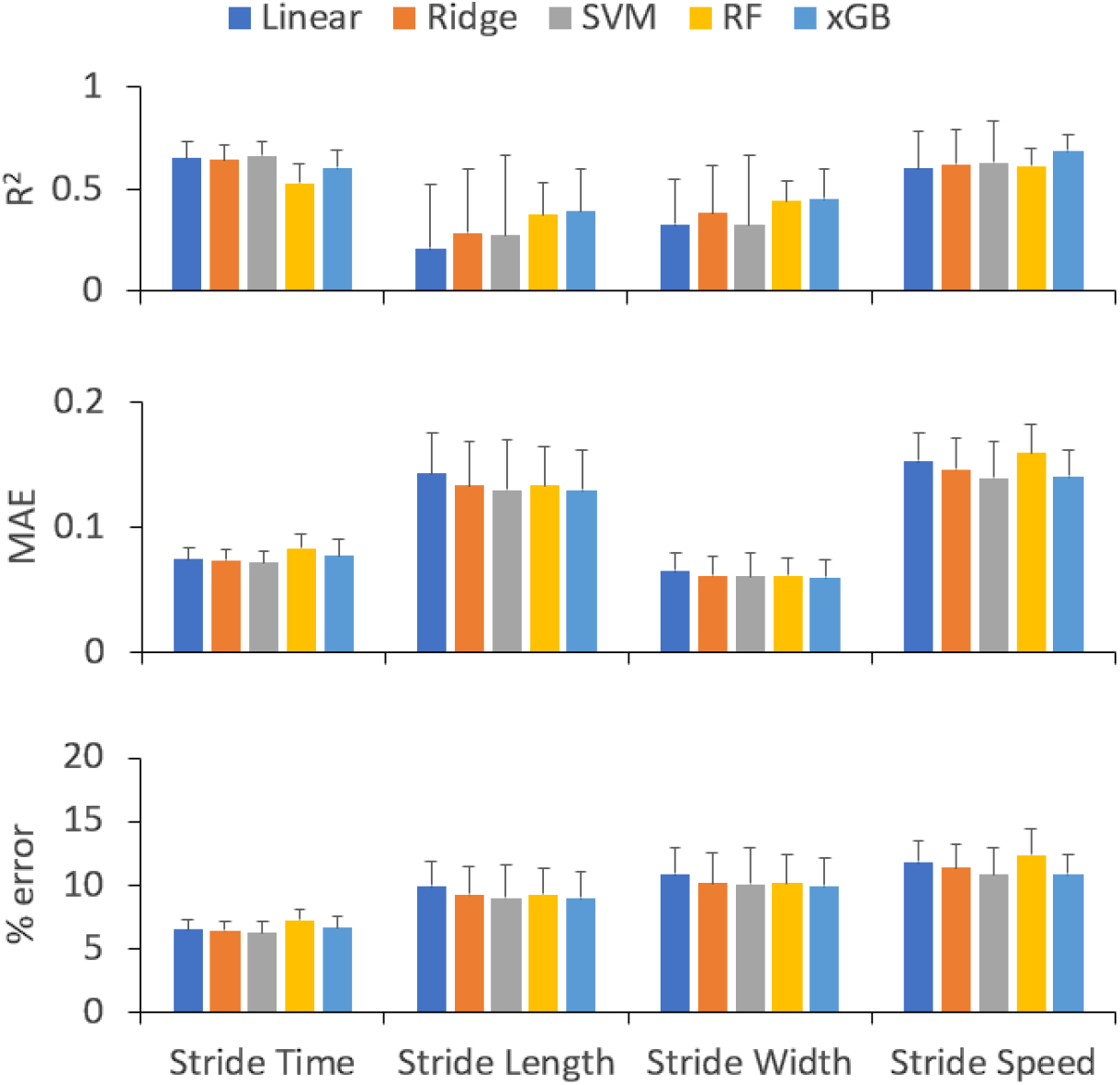
Accuracy of regression models at predicting stride outcomes from smartwatch inertial measurement unit features. Accuracy metrics are the coefficient of determination (R^2^: top), mean absolute error (MAE: middle), and relative error (% error: bottom). MAE is expressed in seconds (stride time), metres (stride length, stride width), and metres/second (stride speed). Models evaluated were linear, ridge, support vector machine (SVM), random forest (RF), and extreme gradient boosted (xGB) regressors.

Spatiotemporal variability was much more accurately predicted by separate, dedicated models (Figure 2). These predictions were generally best using SVM, with R^2^ = 0.35 ± 0.18 and MAE = 0.38 ± 0.08% for stride time CV, R^2^ = 0.52 ± 0.15 and MAE = 0.42 ± 0.13% for stride length CV, R^2^ = 0.42 ± 0.11 and MAE = 0.55 ± 0.16% for stride width CV, and R^2^ = 0.38 ± 0.24 and MAE = 0.28 ± 0.09% for stride speed CV. MAE values corresponded to relative errors of 18–22%.

**Figure 2.**
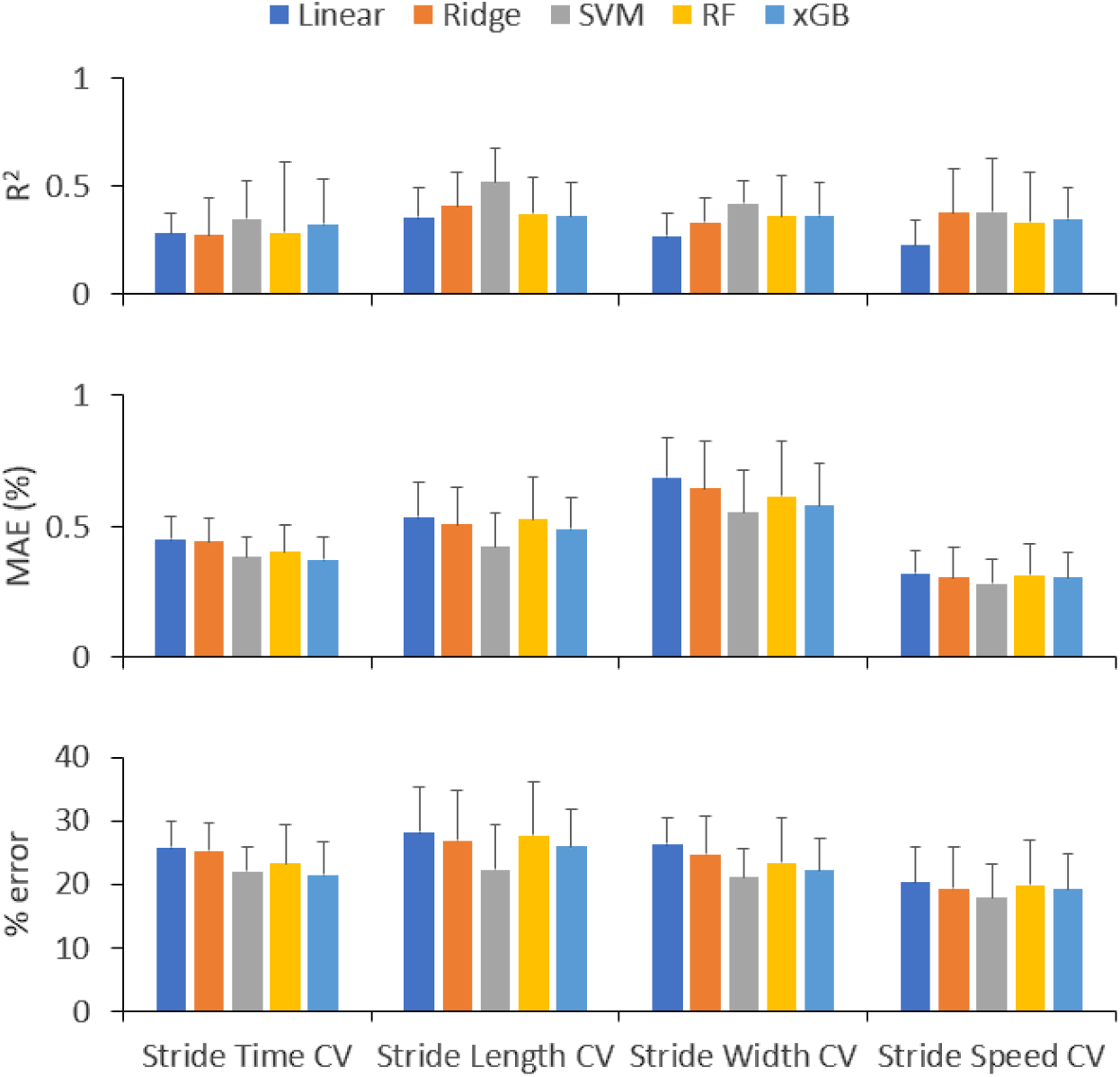
Accuracy of regression models at predicting spatiotemporal variability from smartwatch inertial measurement unit features. Accuracy metrics are the coefficient of determination (R^2^: top), mean absolute error (MAE: middle), and relative error (% error: bottom). MAE is expressed as a percentage. Models evaluated were linear, ridge, support vector machine (SVM), random forest (RF), and extreme gradient boosted (xGB) regressors. CV = coefficient of variation.

### 5.2 Concurrent validity and sensitivity of predictions

Relative to measured values, xGB predictions of stride outcomes had good-to-excellent consistency (ICC_2,1_: 0.60–0.86) and SVM predictions of spatiotemporal variability had fair-to-good consistency (ICC_2,1_: 0.47–0.64) (Table 3). Bland–Altman plots (Figures 3–4) revealed that predictions were not biased on average (i.e. the LOA_95%_ band between the lower and upper limits included zero), although high-variability cases were typically underestimated.

**Table 3.** Consistency (intraclass correlation coefficients [ICC_2,1_]) and agreement (bias, 95% limits of agreement [LOA_95%_]) of smartwatch-based predictions for stride outcomes (using extreme gradient boosting) and for spatiotemporal variability (using support vector machines) relative to optoelectronic-measured values. Spatiotemporal variability was measured by the coefficient of variation (CV).

| Variable | ICC<sub>2,1</sub> | Bias | LOA<sub>95%</sub> |  |
| --- | --- | --- | --- | --- |
|  |  |  | Lower | Upper |
| Stride time (s) | 0.86 | 0.00 | -0.16 | 0.16 |
| Stride length (m) | 0.60 | 0.01 | -0.33 | 0.34 |
| Stride width (m) | 0.60 | 0.00 | -0.15 | 0.16 |
| Stride speed (m/s) | 0.84 | 0.01 | -0.34 | 0.36 |
| Stride time CV (%) | 0.54 | 0.04 | -1.07 | 1.16 |
| Stride length CV (%) | 0.64 | -0.07 | -1.29 | 1.15 |
| Stride width CV (%) | 0.57 | -0.10 | -1.69 | 1.48 |
| Stride speed CV (%) | 0.47 | -0.03 | -0.97 | 0.91 |

**Figure 3.**
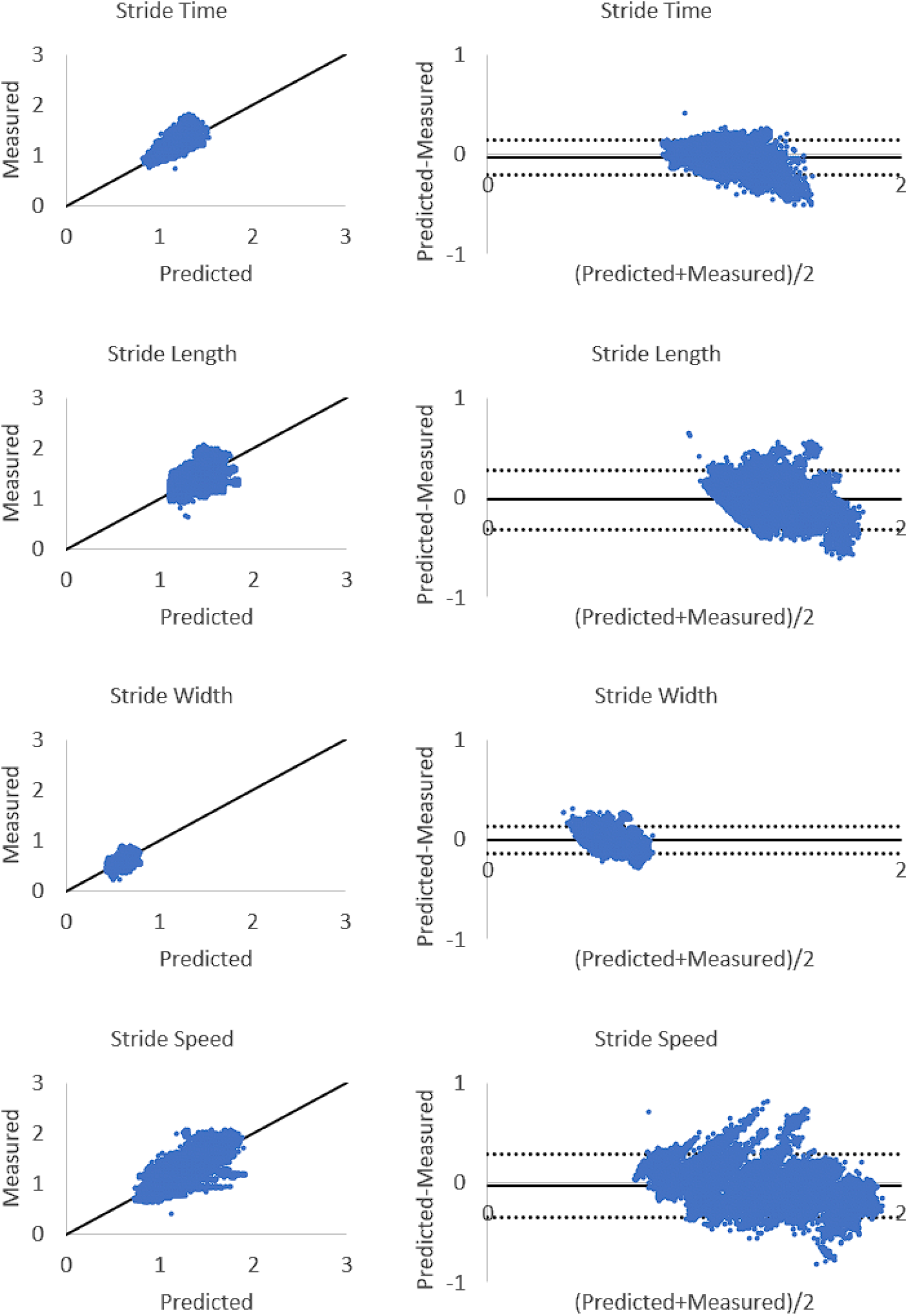
Scatter and Bland–Altman plots of optoelectronic-measured and smartwatch extreme gradient boosting-predicted stride outcomes. Bland–Altman plot lines indicate bias (solid black) and 95% limits of agreement (dotted black).

**Figure 4.**
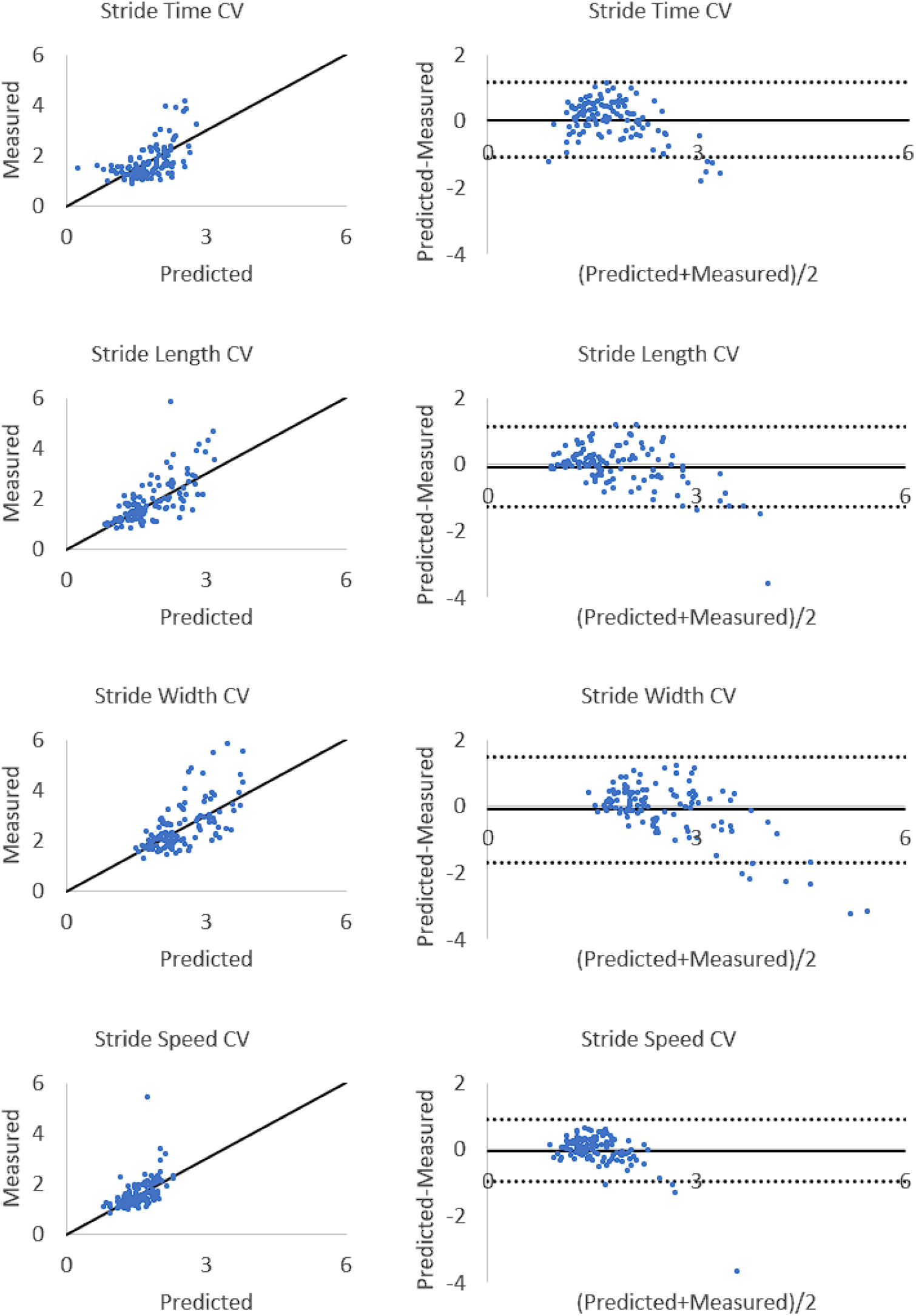
Scatter and Bland–Altman plots of optoelectronic-measured and smartwatch support vector machine-predicted spatiotemporal variability. Bland–Altman plot lines indicate bias (solid black) and 95% limits of agreement (dotted black).

Condition main effects revealed a general pattern of shorter-duration, longer, wider, faster, and less variable strides with increased preferred speed (stride time: p < 0.001, η^2^ = 0.91; stride length: p < 0.001, η^2^ = 0.90; stride width: p < 0.001, η^2^ = 0.90; stride speed: p < 0.001, η^2^ = 0.90; stride time CV: p < 0.001, η^2^ = 0.83; stride length CV: p < 0.001, η^2^ = 0.87; stride width CV: p < 0.001, η^2^ = 0.89; stride speed CV: p < 0.001, η^2^ = 0.80) (Figures 5–6). Magnitude of condition responses were not fully consistent between measured and predicted values, as demonstrated by significant Condition × Model interactions on stride time (p < 0.001, η^2^ = 0.27), stride length (p < 0.001, η^2^ = 0.27), stride width (p < 0.001, η^2^ = 0.25), stride speed (p < 0.001, η^2^ = 0.29), stride time CV (p < 0.001, η^2^ = 0.10), stride length CV (p < 0.001, η^2^ = 0.12), and stride width CV (p < 0.001, η^2^ = 0.14). Condition × Model contrasts of stride time, length, width, and speed were significant for xGB regressors (p < 0.00625), indicating an underestimation of the average predicted speed-related change for stride outcomes. Condition × Model contrasts of stride time CV, stride length CV, and stride width CV were not significant for SVM regressors (p ≥ 0.00625), indicating no over- or underestimation in the average predicted speed-related change for spatiotemporal variability.

**Figure 5.**
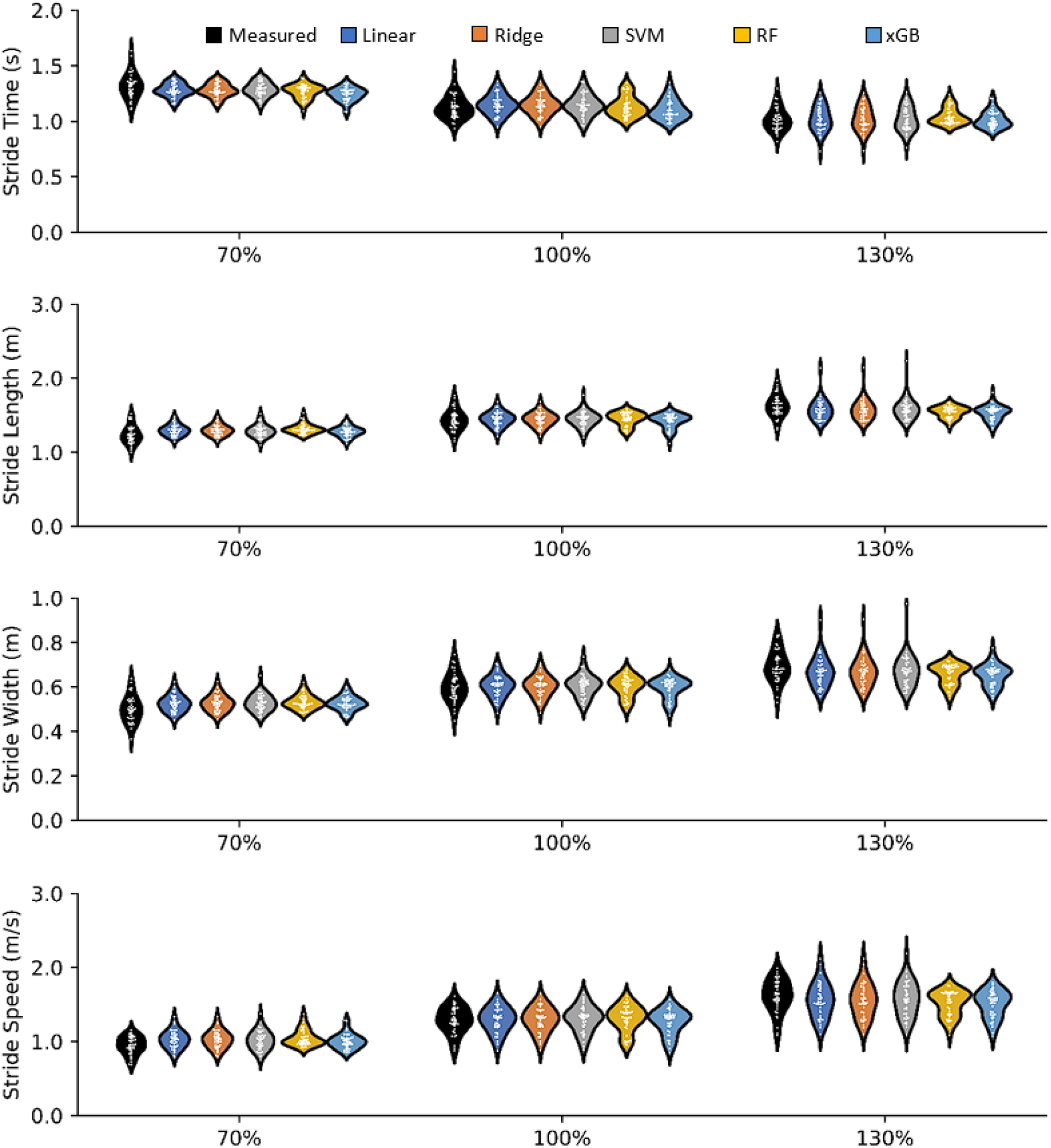
Optoelectronic-measured and smartwatch-based predictions of stride outcomes during gait at 70%, 100%, and 130% of preferred speed. Models evaluated were linear, ridge, support vector machine (SVM), random forest (RF), and extreme gradient boosting (xGB) regressors.

**Figure 6.**
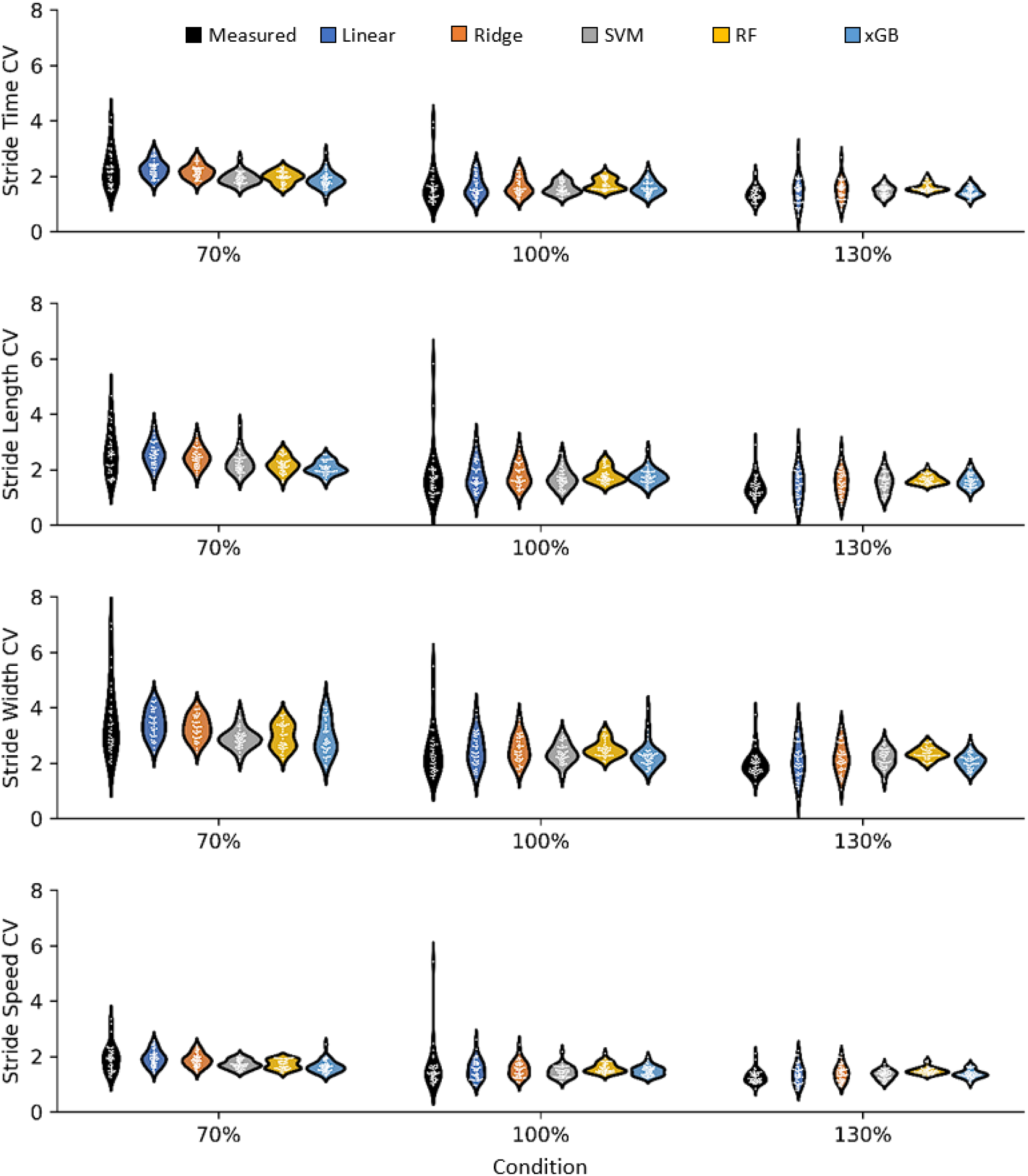
Optoelectronic-measured and smartwatch-based predictions of spatiotemporal variability during gait at 70%, 100%, and 130% of preferred speed. Variability was quantified by the coefficient of variation (CV). Models evaluated were linear, ridge, support vector machine (SVM), random forest (RF), and extreme gradient boosting (xGB) regressors.

## 6. Discussion

Using regression-based machine learning, we developed smartwatch models that explain 39–69% of spatiotemporal stride output and 35–52% of spatiotemporal variability during treadmill walking in healthy young adults. Our main findings pertaining to these models are (1) spatiotemporal variability was predicted more accurately by dedicated regression models than by calculating from single-stride predictions, (2) spatiotemporal predictions had fair-to-excellent consistency and were not biased on average, (3) high spatiotemporal variability cases were typically underestimated, and (4) predicted spatiotemporal responses were sensitive to large within-subject effects (i.e. altered gait speed).

Before modelling spatiotemporal variability, we first evaluated the accuracy of single-stride spatiotemporal outcome predictions by the smartwatch, finding relative errors of 7% for stride time, 9% for stride length, 10% for stride width, and 11% for stride speed. Stride time errors obtained in this study (7%) improve upon the 11% errors in a previous wrist IMU model that was based on peak-to-peak duration of anterior–posterior wrist angular velocity ^25^, and upon the 9–15% errors in step, stance, and swing time from a model of smartwatch IMU features from both wrists, based on means reported in young adults ^5,12^. However, model accuracy was poorer compared to IMU applications on other regions of the body. The most common position reported in the literature is on the foot; using physics-based or machine learning-based approaches, previous foot IMU models measured temporal parameters with approximately 4% error ^42,44,48,49^ and spatial parameters with 1–5% error ^13,15,35,42,44,48,49^. Similar errors of 2% in stride length and speed have been achieved by shank-based IMU models ^27^ and errors of 3–14% in spatiotemporal outputs were reported by trunk and pelvis-based IMU models ^14,36^. Collectively, these findings illustrate a slight improvement in accuracy of single-stride spatiotemporal predictions achieved by our single-smartwatch approach relative to previous wrist IMU and bilateral smartwatch IMU applications, but a lower accuracy relative to IMUs positioned on anatomical segments closer to foot-ground contact. There is a clear trade-off between choosing an IMU position with higher single-stride spatiotemporal accuracy and an IMU position that is preferred by users ^32^ and widely available as a consumer product.

Single-stride spatiotemporal prediction errors were amplified when used to calculate variability (errors of 63–272%) relative to the dedicated models built to predict spatiotemporal variability (errors of 18–22%). The SVM regression models with non-linear kernels outperformed linear and ridge regression, indicating that the relationship between wrist movement patterns and the spatial and temporal variability of foot placement is mostly non-linear. This phenomenon likely explains, in part, why our model predicted stride time CV more accurately (R^2^ = 0.35) than in a previous linear regression model (R^2^ = 0.22) ^1^, with models also differing in the location and the modality of motion inputs (smartwatch-based wrist movement patterns vs. optoelectronic-based lower limb joint motor patterns). Although extreme gradient boosting performed best for single-stride outcomes, this technique likely overfit for spatiotemporal variability, despite our effort to control this risk with an early termination criterion during model training. With 117 total trials, each test set had 23 or 24 trials, compared to the 4600 or 4800 strides in each single-stride test set. Thus, more spatiotemporal variability data are likely needed to prevent overfitting by ensemble techniques.

Smartwatch IMU-based predictions were less accurate for spatiotemporal variability than for single-stride outcomes in all cases. Although the larger relative errors for spatiotemporal variability could be due, in part, to differences in smartwatch IMU Metrics and machine learning model selection, we hypothesize that they reflect a combination of the smaller dataset available for model training and the greater challenge in quantifying stride-to-stride variability compared to average stride spatiotemporal parameters based solely on wrist movement. Similar to single-stride outcomes, spatiotemporal variability is likely to be more accurately predicted by IMUs closer to ground contact. Using a foot IMU, Rebula et al. ^35^ predicted stride length and width variability with relative errors within 4% and good-to-excellent consistency, better than the relative errors of 21–22% and fair-to-good consistency observed in our study using a smartwatch IMU. As with single-stride outcomes, predicting spatiotemporal variability requires one to consider the competing objectives of achieving high accuracy with a foot-based IMU and achieving high user compliance with a wrist-based IMU.

The resolution of this trade-off depends on the application of interest. The sensitivity of predictions to changes in gait speed in this study demonstrate that smartwatch-predicted single-stride and spatiotemporal variability responses during constant-speed walking are sensitive to within-subject phenomena with large effect sizes. As evaluations continue to leave the lab, driven by the need for dozens or upwards of one hundred strides for reliable assessment ^24,37^ and the debate on whether movement patterns during treadmill gait accurately represent those seen overground ^19,20^, our smartwatch-based models provide a new user-preferred method with potential for widespread, real-world application. For example, as a research tool, the smartwatch-based model could be used to investigate how single-stride and spatiotemporal variability gait patterns respond to different terrain, temperatures, and weather conditions. However, we strongly caution readers on its use as a tool to predict spatiotemporal variability for the purposes of fall risk assessment in its present form. On one hand, we demonstrate strong feasibility based on values reported in the literature for older fallers and non-fallers ^26^. The 1.68%, 1.41%, and 2.99% differences in stride time CV, stride length CV, and stride speed CV are larger than the 0.38%, 0.42%, 0.28% absolute errors of our models. Yet, our models are limited to features extracted from young adults, whose arm swing and spatiotemporal gait patterns differ from those of older adults ^20,29^. Further work is needed to refine the smartwatch-based model in a larger sample of adults across the lifespan, including older adults with and without a history of falls.

In conclusion, from top-scoring smartwatch IMU Metrics, we developed SVM models that predicted single-stride spatiotemporal outcomes with 7–11% relative error and xGB models that predicted spatiotemporal variability with 0.28–0.55% absolute error and 18–22% relative error in treadmill gait of young adults. Predictions had fair-to-excellent consistency with optoelectronic-measured values and were sensitive to detecting large effect sizes. Spatiotemporal variability prediction errors were smaller than reported differences between fallers and non-fallers, yet further assessment is needed in these older populations and in larger numbers with our models. For the first time, we demonstrate the feasibility of using a single smartwatch IMU to evaluate spatiotemporal variability in gait, a step towards more widespread real-world continuous monitoring.

## Acknowledgement

The authors thank the participants for volunteering their time.

