## Appendix A and B for "Smartwatch-based prediction of single-stride and stride-to-stride gait outcomes using regression-based machine learning"

**Appendix A: Hyperparameter tuning options for random search cross-validation**

Table A1. Hyperparameter tuning options for each regressor. Hyperparameter details can be found online in the Scikit-learn documentation for Python (<https://scikit-learn.org/stable/>).

| Regressor | Hyperparameter | Hyperparameter options |
| --- | --- | --- |
| Ridge | solver | [auto, svd, cholesky, lsqr, sparse_cg] |
|  | alpha | [1e-15, 1e-10, 1e-8, 1e-4, 1e-3, 1e-2, 1, 5, 10, 20] |
| Support vector | kernel | [linear, rbf, sigmoid, poly] |
|  | gamma | [0.001, 0.001, 0.01, 0.1, 1, 10] |
|  | C | [0.1, 0.5, 1, 5, 10, 50, 100] |
| Random forest | n_estimators | [50, 100, 200] |
|  | min_samples_split | [0.1, 0.3, 0.5] |
|  | min_samples_leaf | [0.1, 0.3, 0.5] |
|  | max_leaf_nodes | [2, 3, 4, 5, 6, 7] |
|  | max_features | [1, 2, 3, 4, 5, 6, 7, 8, 9, 10, 11, 12, 13, 14, 15, 16, 17, 18, 19, 20, 21, 22, 23, 24] |
|  | max_depth | [1, 2, 3, 4, 5] |
|  | bootstrap | [True, False] |
| Extreme gradient boosting | n_estimators | [50, 100, 200] |
|  | max_depth | [1, 2, 3, 4, 5] |
|  | learning_rate | [0.005, 0.01, 0.015, 0.02] |
|  | early_stopping_rounds | [10] |

**Appendix B: Hyperparameter-tuned regression models**

Table B1. Final hyperparameters for each variable, regressor, and repeated fold. Hyperparameter details can be found online in the Scikit-learn documentation for Python (<https://scikit-learn.org/stable/>).

|  | Regressor | Outer fold | Hyperparameters |
| --- | --- | --- | --- |
| Stride time (s) | Ridge | 1 | {'solver': 'sparse_cg', 'alpha': 20} |
|  |  | 2 | {'solver': 'lsqr', 'alpha': 1e-15} |
|  |  | 3 | {'solver': 'lsqr', 'alpha': 5} |
|  |  | 4 | {'solver': 'sparse_cg', 'alpha': 10} |
|  |  | 5 | {'solver': 'sparse_cg', 'alpha': 10} |
|  | Support vector | 1 | {'kernel': 'linear', 'gamma': 0.0001, 'C': 100} |
|  |  | 2 | {'kernel': 'linear', 'gamma': 0.0001, 'C': 100} |
|  |  | 3 | {'kernel': 'linear', 'gamma': 0.0001, 'C': 0.1} |
|  |  | 4 | {'kernel': 'linear', 'gamma': 0.0001, 'C': 100} |
|  |  | 5 | {'kernel': 'linear', 'gamma': 0.0001, 'C': 5} |
|  | Random forest | 1 | {'n_estimators': 200, 'min_samples_split': 0.1, 'min_samples_leaf': 0.1, 'max_leaf_nodes': 7, 'max_features': 8, 'max_depth': 3, 'bootstrap': True} |
|  |  | 2 | {'n_estimators': 100, 'min_samples_split': 0.1, 'min_samples_leaf': 0.1, 'max_leaf_nodes': 7, 'max_features': 9, 'max_depth': 5, 'bootstrap': False} |
|  |  | 3 | {'n_estimators': 200, 'min_samples_split': 0.1, 'min_samples_leaf': 0.1, 'max_leaf_nodes': 7, 'max_features': 8, 'max_depth': 3, 'bootstrap': True} |
|  |  | 4 | {'n_estimators': 100, 'min_samples_split': 0.1, 'min_samples_leaf': 0.1, 'max_leaf_nodes': 7, 'max_features': 9, 'max_depth': 5, 'bootstrap': False} |
|  |  | 5 | {'n_estimators': 50, 'min_samples_split': 0.1, 'min_samples_leaf': 0.1, 'max_leaf_nodes': 7, 'max_features': 8, 'max_depth': 4, 'bootstrap': True} |
|  | Extreme gradient boosting | 1 | {'n_estimators': 200, 'max_depth': 5, 'learning_rate': 0.02, 'early_stopping_rounds': 10} |
|  |  | 2 | {'n_estimators': 200, 'max_depth': 5, 'learning_rate': 0.02, 'early_stopping_rounds': 10} |
|  |  | 3 | {'n_estimators': 200, 'max_depth': 5, 'learning_rate': 0.015, 'early_stopping_rounds': 10} |
|  |  | 4 | {'n_estimators': 200, 'max_depth': 5, 'learning_rate': 0.02, 'early_stopping_rounds': 10} |
|  |  | 5 | {'n_estimators': 200, 'max_depth': 5, 'learning_rate': 0.02, 'early_stopping_rounds': 10} |
| Stride length (m) | Ridge | 1 | {'solver': 'lsqr', 'alpha': 20} |
|  |  | 2 | {'solver': 'auto', 'alpha': 20} |
|  |  | 3 | {'solver': 'auto', 'alpha': 20} |
|  |  | 4 | {'solver': 'lsqr', 'alpha': 20} |
|  |  | 5 | {'solver': 'lsqr', 'alpha': 5} |
|  | Support vector | 1 | {'kernel': 'linear', 'gamma': 0.0001, 'C': 0.1} |
|  |  | 2 | {'kernel': 'linear', 'gamma': 0.0001, 'C': 0.1} |
|  |  | 3 | {'kernel': 'linear', 'gamma': 0.0001, 'C': 0.1} |
|  |  | 4 | {'kernel': 'linear', 'gamma': 0.0001, 'C': 0.1} |
|  |  | 5 | {'kernel': 'linear', 'gamma': 0.0001, 'C': 0.1} |
|  | Random forest | 1 | {'n_estimators': 50, 'min_samples_split': 0.3, 'min_samples_leaf': 0.1, 'max_leaf_nodes': 4, 'max_features': 9, 'max_depth': 2, 'bootstrap': False} |
|  |  | 2 | {'n_estimators': 50, 'min_samples_split': 0.1, 'min_samples_leaf': 0.1, 'max_leaf_nodes': 7, 'max_features': 8, 'max_depth': 4, 'bootstrap': True} |
|  |  | 3 | {'n_estimators': 50, 'min_samples_split': 0.1, 'min_samples_leaf': 0.1, 'max_leaf_nodes': 7, 'max_features': 8, 'max_depth': 4, 'bootstrap': True} |
|  |  | 4 | {'n_estimators': 200, 'min_samples_split': 0.5, 'min_samples_leaf': 0.1, 'max_leaf_nodes': 2, 'max_features': 13, 'max_depth': 2, 'bootstrap': True} |
|  |  | 5 | {'n_estimators': 100, 'min_samples_split': 0.1, 'min_samples_leaf': 0.1, 'max_leaf_nodes': 7, 'max_features': 9, 'max_depth': 5, 'bootstrap': False} |
|  | Extreme gradient boosting | 1 | {'n_estimators': 200, 'max_depth': 1, 'learning_rate': 0.02, 'early_stopping_rounds': 10} |
|  |  | 2 | {'n_estimators': 200, 'max_depth': 1, 'learning_rate': 0.02, 'early_stopping_rounds': 10} |
|  |  | 3 | {'n_estimators': 200, 'max_depth': 4, 'learning_rate': 0.02, 'early_stopping_rounds': 10} |
|  |  | 4 | {'n_estimators': 200, 'max_depth': 4, 'learning_rate': 0.02, 'early_stopping_rounds': 10} |
|  |  | 5 | {'n_estimators': 200, 'max_depth': 4, 'learning_rate': 0.02, 'early_stopping_rounds': 10} |
| Stride width (m) | Ridge | 1 | {'solver': 'lsqr', 'alpha': 20} |
|  |  | 2 | {'solver': 'sparse_cg', 'alpha': 20} |
|  |  | 3 | {'solver': 'auto', 'alpha': 20} |
|  |  | 4 | {'solver': 'lsqr', 'alpha': 20} |
|  |  | 5 | {'solver': 'lsqr', 'alpha': 5} |
|  | Support vector | 1 | {'kernel': 'linear', 'gamma': 0.0001, 'C': 0.1} |
|  |  | 2 | {'kernel': 'linear', 'gamma': 0.0001, 'C': 5} |
|  |  | 3 | {'kernel': 'linear', 'gamma': 0.0001, 'C': 5} |
|  |  | 4 | {'kernel': 'linear', 'gamma': 0.0001, 'C': 0.1} |
|  |  | 5 | {'kernel': 'linear', 'gamma': 0.0001, 'C': 0.1} |
|  | Random forest | 1 | {'n_estimators': 200, 'min_samples_split': 0.1, 'min_samples_leaf': 0.1, 'max_leaf_nodes': 4, 'max_features': 13, 'max_depth': 2, 'bootstrap': False} |
|  |  | 2 | {'n_estimators': 50, 'min_samples_split': 0.3, 'min_samples_leaf': 0.1, 'max_leaf_nodes': 6, 'max_features': 18, 'max_depth': 5, 'bootstrap': False} |
|  |  | 3 | {'n_estimators': 100, 'min_samples_split': 0.1, 'min_samples_leaf': 0.1, 'max_leaf_nodes': 7, 'max_features': 9, 'max_depth': 5, 'bootstrap': False} |
|  |  | 4 | {'n_estimators': 200, 'min_samples_split': 0.1, 'min_samples_leaf': 0.1, 'max_leaf_nodes': 3, 'max_features': 20, 'max_depth': 5, 'bootstrap': True} |
|  |  | 5 | {'n_estimators': 100, 'min_samples_split': 0.1, 'min_samples_leaf': 0.1, 'max_leaf_nodes': 7, 'max_features': 9, 'max_depth': 5, 'bootstrap': False} |
|  | Extreme gradient boosting | 1 | {'n_estimators': 200, 'max_depth': 1, 'learning_rate': 0.015, 'early_stopping_rounds': 10} |
|  |  | 2 | {'n_estimators': 200, 'max_depth': 1, 'learning_rate': 0.015, 'early_stopping_rounds': 10} |
|  |  | 3 | {'n_estimators': 200, 'max_depth': 5, 'learning_rate': 0.02, 'early_stopping_rounds': 10} |
|  |  | 4 | {'n_estimators': 200, 'max_depth': 4, 'learning_rate': 0.015, 'early_stopping_rounds': 10} |
|  |  | 5 | {'n_estimators': 200, 'max_depth': 4, 'learning_rate': 0.02, 'early_stopping_rounds': 10} |
| Stride speed (m/s) | Ridge | 1 | {'solver': 'sparse_cg', 'alpha': 20} |
|  |  | 2 | {'solver': 'lsqr', 'alpha': 20} |
|  |  | 3 | {'solver': 'lsqr', 'alpha': 1e-15} |
|  |  | 4 | {'solver': 'lsqr', 'alpha': 20} |
|  |  | 5 | {'solver': 'lsqr', 'alpha': 5} |
|  | Support vector | 1 | {'kernel': 'linear', 'gamma': 0.0001, 'C': 0.5} |
|  |  | 2 | {'kernel': 'linear', 'gamma': 0.0001, 'C': 0.1} |
|  |  | 3 | {'kernel': 'linear', 'gamma': 0.0001, 'C': 0.5} |
|  |  | 4 | {'kernel': 'linear', 'gamma': 0.0001, 'C': 0.1} |
|  |  | 5 | {'kernel': 'linear', 'gamma': 0.0001, 'C': 100} |
|  | Random forest | 1 | {'n_estimators': 50, 'min_samples_split': 0.3, 'min_samples_leaf': 0.1, 'max_leaf_nodes': 6, 'max_features': 18, 'max_depth': 5, 'bootstrap': False} |
|  |  | 2 | {'n_estimators': 50, 'min_samples_split': 0.3, 'min_samples_leaf': 0.1, 'max_leaf_nodes': 6, 'max_features': 18, 'max_depth': 5, 'bootstrap': False} |
|  |  | 3 | {'n_estimators': 100, 'min_samples_split': 0.1, 'min_samples_leaf': 0.1, 'max_leaf_nodes': 7, 'max_features': 9, 'max_depth': 5, 'bootstrap': False} |
|  |  | 4 | {'n_estimators': 200, 'min_samples_split': 0.1, 'min_samples_leaf': 0.1, 'max_leaf_nodes': 4, 'max_features': 13, 'max_depth': 2, 'bootstrap': False} |
|  |  | 5 | {'n_estimators': 100, 'min_samples_split': 0.1, 'min_samples_leaf': 0.1, 'max_leaf_nodes': 7, 'max_features': 9, 'max_depth': 5, 'bootstrap': False} |
|  | Extreme gradient boosting | 1 | {'n_estimators': 200, 'max_depth': 5, 'learning_rate': 0.02, 'early_stopping_rounds': 10} |
|  |  | 2 | {'n_estimators': 200, 'max_depth': 5, 'learning_rate': 0.02, 'early_stopping_rounds': 10} |
|  |  | 3 | {'n_estimators': 200, 'max_depth': 5, 'learning_rate': 0.02, 'early_stopping_rounds': 10} |
|  |  | 4 | {'n_estimators': 200, 'max_depth': 5, 'learning_rate': 0.02, 'early_stopping_rounds': 10} |
|  |  | 5 | {'n_estimators': 200, 'max_depth': 5, 'learning_rate': 0.02, 'early_stopping_rounds': 10} |
| Stride time CV (%) | Ridge | 1 | {'solver': 'lsqr', 'alpha': 20} |
|  |  | 2 | {'solver': 'sparse_cg', 'alpha': 20} |
|  |  | 3 | {'solver': 'sparse_cg', 'alpha': 20} |
|  |  | 4 | {'solver': 'lsqr', 'alpha': 0.01} |
|  |  | 5 | {'solver': 'auto', 'alpha': 1} |
|  | Support vector | 1 | {'kernel': 'rbf', 'gamma': 0.01, 'C': 5} |
|  |  | 2 | {'kernel': 'rbf', 'gamma': 0.01, 'C': 10} |
|  |  | 3 | {'kernel': 'rbf', 'gamma': 0.001, 'C': 100} |
|  |  | 4 | {'kernel': 'rbf', 'gamma': 0.1, 'C': 5} |
|  |  | 5 | {'kernel': 'rbf', 'gamma': 0.01, 'C': 10} |
|  | Random forest | 1 | {'n_estimators': 50, 'min_samples_split': 0.5, 'min_samples_leaf': 0.1, 'max_leaf_nodes': 4, 'max_features': 2, 'max_depth': 2, 'bootstrap': True} |
|  |  | 2 | {'n_estimators': 200, 'min_samples_split': 0.3, 'min_samples_leaf': 0.1, 'max_leaf_nodes': 3, 'max_features': 17, 'max_depth': 3, 'bootstrap': False} |
|  |  | 3 | {'n_estimators': 50, 'min_samples_split': 0.5, 'min_samples_leaf': 0.3, 'max_leaf_nodes': 2, 'max_features': 3, 'max_depth': 1, 'bootstrap': False} |
|  |  | 4 | {'n_estimators': 50, 'min_samples_split': 0.5, 'min_samples_leaf': 0.3, 'max_leaf_nodes': 2, 'max_features': 3, 'max_depth': 1, 'bootstrap': False} |
|  |  | 5 | {'n_estimators': 200, 'min_samples_split': 0.5, 'min_samples_leaf': 0.3, 'max_leaf_nodes': 5, 'max_features': 5, 'max_depth': 1, 'bootstrap': False} |
|  | Extreme gradient boosting | 1 | {'n_estimators': 200, 'max_depth': 1, 'learning_rate': 0.02, 'early_stopping_rounds': 10} |
|  |  | 2 | {'n_estimators': 200, 'max_depth': 1, 'learning_rate': 0.015, 'early_stopping_rounds': 10} |
|  |  | 3 | {'n_estimators': 200, 'max_depth': 1, 'learning_rate': 0.02, 'early_stopping_rounds': 10} |
|  |  | 4 | {'n_estimators': 200, 'max_depth': 1, 'learning_rate': 0.02, 'early_stopping_rounds': 10} |
|  |  | 5 | {'n_estimators': 200, 'max_depth': 2, 'learning_rate': 0.02, 'early_stopping_rounds': 10} |
| Stride length CV (%) | Ridge | 1 | {'solver': 'lsqr', 'alpha': 20} |
|  |  | 2 | {'solver': 'lsqr', 'alpha': 10} |
|  |  | 3 | {'solver': 'sparse_cg', 'alpha': 5} |
|  |  | 4 | {'solver': 'sparse_cg', 'alpha': 1} |
|  |  | 5 | {'solver': 'lsqr', 'alpha': 10} |
|  | Support vector | 1 | {'kernel': 'rbf', 'gamma': 0.01, 'C': 10} |
|  |  | 2 | {'kernel': 'rbf', 'gamma': 0.01, 'C': 10} |
|  |  | 3 | {'kernel': 'rbf', 'gamma': 0.01, 'C': 10} |
|  |  | 4 | {'kernel': 'rbf', 'gamma': 0.01, 'C': 10} |
|  |  | 5 | {'kernel': 'rbf', 'gamma': 0.01, 'C': 10} |
|  | Random forest | 1 | {'n_estimators': 200, 'min_samples_split': 0.1, 'min_samples_leaf': 0.1, 'max_leaf_nodes': 7, 'max_features': 8, 'max_depth': 3, 'bootstrap': True} |
|  |  | 2 | {'n_estimators': 50, 'min_samples_split': 0.3, 'min_samples_leaf': 0.1, 'max_leaf_nodes': 5, 'max_features': 2, 'max_depth': 3, 'bootstrap': False} |
|  |  | 3 | {'n_estimators': 50, 'min_samples_split': 0.1, 'min_samples_leaf': 0.1, 'max_leaf_nodes': 7, 'max_features': 8, 'max_depth': 4, 'bootstrap': True} |
|  |  | 4 | {'n_estimators': 200, 'min_samples_split': 0.5, 'min_samples_leaf': 0.1, 'max_leaf_nodes': 3, 'max_features': 17, 'max_depth': 5, 'bootstrap': False} |
|  |  | 5 | {'n_estimators': 200, 'min_samples_split': 0.1, 'min_samples_leaf': 0.1, 'max_leaf_nodes': 7, 'max_features': 8, 'max_depth': 3, 'bootstrap': True} |
|  | Extreme gradient boosting | 1 | {'n_estimators': 200, 'max_depth': 1, 'learning_rate': 0.015, 'early_stopping_rounds': 10} |
|  |  | 2 | {'n_estimators': 200, 'max_depth': 1, 'learning_rate': 0.01, 'early_stopping_rounds': 10} |
|  |  | 3 | {'n_estimators': 200, 'max_depth': 1, 'learning_rate': 0.02, 'early_stopping_rounds': 10} |
|  |  | 4 | {'n_estimators': 200, 'max_depth': 1, 'learning_rate': 0.02, 'early_stopping_rounds': 10} |
|  |  | 5 | {'n_estimators': 200, 'max_depth': 1, 'learning_rate': 0.02, 'early_stopping_rounds': 10} |
| Stride width CV (%) | Ridge | 1 | {'solver': 'sparse_cg', 'alpha': 20} |
|  |  | 2 | {'solver': 'sparse_cg', 'alpha': 20} |
|  |  | 3 | {'solver': 'auto', 'alpha': 20} |
|  |  | 4 | {'solver': 'lsqr', 'alpha': 1} |
|  |  | 5 | {'solver': 'sparse_cg', 'alpha': 20} |
|  | Support vector | 1 | {'kernel': 'rbf', 'gamma': 0.01, 'C': 10} |
|  |  | 2 | {'kernel': 'rbf', 'gamma': 0.01, 'C': 5} |
|  |  | 3 | {'kernel': 'rbf', 'gamma': 0.01, 'C': 5} |
|  |  | 4 | {'kernel': 'rbf', 'gamma': 0.01, 'C': 5} |
|  |  | 5 | {'kernel': 'rbf', 'gamma': 0.01, 'C': 10} |
|  | Random forest | 1 | {'n_estimators': 50, 'min_samples_split': 0.3, 'min_samples_leaf': 0.1, 'max_leaf_nodes': 5, 'max_features': 2, 'max_depth': 3, 'bootstrap': False} |
|  |  | 2 | {'n_estimators': 50, 'min_samples_split': 0.3, 'min_samples_leaf': 0.1, 'max_leaf_nodes': 5, 'max_features': 2, 'max_depth': 3, 'bootstrap': False} |
|  |  | 3 | {'n_estimators': 50, 'min_samples_split': 0.3, 'min_samples_leaf': 0.1, 'max_leaf_nodes': 5, 'max_features': 2, 'max_depth': 3, 'bootstrap': False} |
|  |  | 4 | {'n_estimators': 50, 'min_samples_split': 0.1, 'min_samples_leaf': 0.1, 'max_leaf_nodes': 7, 'max_features': 8, 'max_depth': 4, 'bootstrap': True} |
|  |  | 5 | {'n_estimators': 50, 'min_samples_split': 0.3, 'min_samples_leaf': 0.1, 'max_leaf_nodes': 5, 'max_features': 2, 'max_depth': 3, 'bootstrap': False} |
|  | Extreme gradient boosting | 1 | {'n_estimators': 200, 'max_depth': 1, 'learning_rate': 0.015, 'early_stopping_rounds': 10} |
|  |  | 2 | {'n_estimators': 200, 'max_depth': 1, 'learning_rate': 0.02, 'early_stopping_rounds': 10} |
|  |  | 3 | {'n_estimators': 200, 'max_depth': 1, 'learning_rate': 0.02, 'early_stopping_rounds': 10} |
|  |  | 4 | {'n_estimators': 200, 'max_depth': 2, 'learning_rate': 0.02, 'early_stopping_rounds': 10} |
|  |  | 5 | {'n_estimators': 200, 'max_depth': 1, 'learning_rate': 0.02, 'early_stopping_rounds': 10} |
| Stride speed CV (%) | Ridge | 1 | {'solver': 'lsqr', 'alpha': 20} |
|  |  | 2 | {'solver': 'lsqr', 'alpha': 10} |
|  |  | 3 | {'solver': 'auto', 'alpha': 10} |
|  |  | 4 | {'solver': 'auto', 'alpha': 10} |
|  |  | 5 | {'solver': 'lsqr', 'alpha': 10} |
|  | Support vector | 1 | {'kernel': 'rbf', 'gamma': 0.01, 'C': 10} |
|  |  | 2 | {'kernel': 'rbf', 'gamma': 0.01, 'C': 5} |
|  |  | 3 | {'kernel': 'rbf', 'gamma': 0.001, 'C': 100} |
|  |  | 4 | {'kernel': 'linear', 'gamma': 0.01, 'C': 1} |
|  |  | 5 | {'kernel': 'rbf', 'gamma': 0.01, 'C': 10} |
|  | Random forest | 1 | {'n_estimators': 50, 'min_samples_split': 0.3, 'min_samples_leaf': 0.1, 'max_leaf_nodes': 5, 'max_features': 2, 'max_depth': 3, 'bootstrap': False} |
|  |  | 2 | {'n_estimators': 100, 'min_samples_split': 0.1, 'min_samples_leaf': 0.1, 'max_leaf_nodes': 7, 'max_features': 9, 'max_depth': 5, 'bootstrap': False} |
|  |  | 3 | {'n_estimators': 200, 'min_samples_split': 0.1, 'min_samples_leaf': 0.1, 'max_leaf_nodes': 7, 'max_features': 8, 'max_depth': 3, 'bootstrap': True} |
|  |  | 4 | {'n_estimators': 50, 'min_samples_split': 0.1, 'min_samples_leaf': 0.1, 'max_leaf_nodes': 7, 'max_features': 8, 'max_depth': 4, 'bootstrap': True} |
|  |  | 5 | {'n_estimators': 200, 'min_samples_split': 0.5, 'min_samples_leaf': 0.3, 'max_leaf_nodes': 4, 'max_features': 3, 'max_depth': 5, 'bootstrap': False} |
|  | Extreme gradient boosting | 1 | {'n_estimators': 200, 'max_depth': 1, 'learning_rate': 0.015, 'early_stopping_rounds': 10} |
|  |  | 2 | {'n_estimators': 200, 'max_depth': 2, 'learning_rate': 0.02, 'early_stopping_rounds': 10} |
|  |  | 3 | {'n_estimators': 200, 'max_depth': 4, 'learning_rate': 0.015, 'early_stopping_rounds': 10} |
|  |  | 4 | {'n_estimators': 200, 'max_depth': 1, 'learning_rate': 0.02, 'early_stopping_rounds': 10} |
|  |  | 5 | {'n_estimators': 200, 'max_depth': 1, 'learning_rate': 0.02, 'early_stopping_rounds': 10} |
